# Graph-Based Modeling of Alzheimer’s Protein Interactions via Spiking Neural, Hyperdimensional Encoding, and Scalable Ray-Based Learning

**DOI:** 10.1101/2025.04.11.647919

**Authors:** Saba Zare

## Abstract

This study introduces a novel computational framework for predicting protein-protein interactions (PPIs) in Alzheimer’s disease by integrating biologically inspired and graph-based learning paradigms. The pipeline combines genetic algorithm-driven feature selection, hyperdimensional encoding for robust semantic representation, and spiking neural networks to capture temporal dynamics. These representations are fused with graph neural network embeddings and processed via scalable nearest-neighbor inference. Unlike conventional models, this multi-view architecture enables biologically grounded prediction under data sparsity, offering interpretable insights into disease-relevant molecular interactions.

## I. Introduction

Protein–protein interactions (PPIs) form the backbone of cellular function and provide essential insight into the molecular mechanisms that drive complex diseases such as Alzheimer’s disease [1]. Available PPI data are often sparse, noisy, and incomplete, while molecular relationships underlying neurodegeneration are highly nonlinear and context dependent. These factors limit the effectiveness of traditional sequence-based and pairwise statistical models and create a pressing need for methods that explicitly exploit network topology, multimodal biological signals, and robust representation strategies [2], [3].

Recent developments in network science and machine learning enable modeling that captures both topological context and learned relational patterns across large biological graphs [4], [5]. Graph Neural Networks can learn structural motifs and propagate information across interactome neighborhoods, but performance may degrade when experimental coverage is limited or when informative features are rare. Complementary representation paradigms such as hyperdimensional encoding and biologically inspired neural simulations offer alternative, noise-tolerant encodings that preserve higher-order relationships and temporal dynamics [6], [7]. Evolutionary search can further identify compact, high-value feature subsets that maximize predictive signal while reducing overfitting risk in low-data regimes [8].

This study presents a unified computational pipeline for prediction of missing PPIs in the context of Alzheimer’s disease. The pipeline integrates graph-aware feature engineering, evolutionary selection, hyperdimensional augmen-tation, neuromorphic embedding via Leaky Integrate and Fire dynamics, and multi-view fusion with Graph Neural Network descriptors. Each component contributes distinct advantages, and their integration enables capture of complementary biological signals including structural, temporal, and topological features. To address concerns regarding biological relevance and interpretability, semantic structure preservation in hyperdimensional projections is empirically evaluated, and spike-based embeddings are interpreted in relation to protein activity and disease mechanisms.

### Key contributions

1. Construction of an Alzheimer’s-specific interaction network through systematic extraction and filtering of public PPI resources, along with biologically informed topological descriptors.
2. Graph-aware feature selection using a genetic algorithm that preserves discriminative biological and structural signals while reducing dimensionality and noise.
3. Application of hyperdimensional encoding to generate robust, high-capacity representations, with empirical validation of semantic preservation in sparse biological graphs.
4. Generation of spike-based embeddings using a Leaky Integrate and Fire spiking neural network to capture temporal and event-driven aspects of protein interactions.
5. Multi-view fusion of spike embeddings, graph descriptors, and GNN representations, followed by nearest-neighbor inference to predict uncharacterized PPIs.
6. Comprehensive empirical evaluation demonstrating improved predictive accuracy, resilience under data sparsity, and interpretability for experimental prioritization.

While each module has demonstrated individual utility,their integration enables a biologically grounded, multi-perspective modeling of protein–protein interactions. This fusion captures complementary signals: structural motifs from graph topology, dynamic regulation from spike-based encoding, and distributed semantic structure from hyper-dimensional projections. Together, these components allow the framework to model nonlinear dependencies and sparse interactions more effectively than conventional architectures, particularly in the context of Alzheimer’s disease where data incompleteness and biological heterogeneity are prevalent.

The remainder of this paper is organized as follows: Section II reviews related work on protein–protein interaction (PPI) prediction and network-based modeling. Section III presents the proposed methodology. Section IV details the experimental results. Section V provides a discussion along with directions for future research, and Section VI concludes the paper.

## II. Related Work

Protein–protein interaction (PPI) prediction has evolved considerably with the rise of computational methods. Early approaches primarily used statistical and sequence-based analyses [1], [2], but were limited by data sparsity and noise, prompting a shift toward more robust machine learning models.

Recent deep learning methods have significantly improved PPI prediction. Architectures such as DeepPPI [9] and other convolutional models [10] have shown success in capturing complex patterns in protein sequences and structures. Meanwhile, graph neural networks (GNNs) have emerged as a powerful tool for learning from biological networks. Models like GCN [4], along with comprehensive surveys [5], [11], highlight the effectiveness of GNNs in capturing topological and relational information in PPI graphs.

Hyperdimensional computing (HDC) is another emerging paradigm, inspired by brain-like distributed representations. Foundational work by Kanerva [6] and applications in bioinformatics [12] have demonstrated HDC’s robustness to noise and suitability for biological data. Recent advances by Verges et al. [13] further show its utility in molecular graph classification, balancing computational efficiency and predictive accuracy.

Spiking neural networks (SNNs) offer biologically plausible models by incorporating time dynamics and event-based processing, with LIF-based models [7], [14] gaining traction. Xiao et al. [15] applied SNNs to graph reasoning, demonstrating energy-efficient performance on structured data like knowledge graphs.

This work builds on these advances by integrating deep learning, graph theory, HDC, and SNNs into a unified framework for PPI prediction in Alzheimer’s disease.

## III. Proposed Methodology

The proposed framework integrates multiple computational techniques to predict missing protein–protein interactions (PPIs) in Alzheimer’s disease.

### A. Data Preprocessing and Supplementary Datasets

PPI data is first extracted from public databases, particularly the BioGRID Alzheimer’s Disease Project [16], and rigorously filtered to retain human proteins and Alzheimer’s-related interactions. The initial feature matrix, denoted by *X* ∈*ℝ* ^*m×d*^ (where *m* is the number of samples and *d* the number of features), is normalized as:

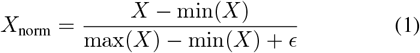

with *ϵ* being a small constant to avoid division by zero. Dimensionality reduction is then performed (e.g., using Singular Value Decomposition) to obtain a compact representation:

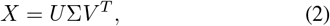

where *U*, Σ, and *V* are the matrices from SVD.

A structured preprocessing pipeline refines the dataset by separating numeric and categorical features. Numeric features are median-imputed, variance-filtered, and standardized. Categorical features undergo constant imputation and one-hot encoding, with high-cardinality values grouped under a generic label.

A column transformer unifies these steps, followed by optional dimensionality reduction using truncated SVD. For high-dimensional inputs, feature selection via variance and mutual information scores retains the most informative attributes.

To enhance protein–protein interaction prediction, three supplementary datasets are incorporated: (i) sequence similarity for evolutionary patterns based on data from UniProt [17], (ii) functional similarity for shared pathways derived from the KEGG database [18], and (iii) structural similarity for 3D conformational insights provided by the RCSB Protein Data Bank [19]. These enrich the biological context of Alzheimer’s-related protein interactions.

### B. Graph-Aware Feature Selection via Genetic Algorithms

A genetic algorithm (GA), implemented using the DEAP framework [20] and enhanced with graph-aware modifications as described in [21], is employed for feature selection on the graph-based dataset. In this setting, a binary vector **b** ∈ {0, 1} ^*n*^ represents the selection of features, where *n* denotes the total number of candidate features. The GA leverages graph-aware genetic operators—crossover, mutation, and fitness evaluation—that account for structural characteristics of the protein-protein interaction (PPI) network, such as node centrality and connectivity.

The fitness evaluation incorporates three main components:

1. **Classifier Performance:** The cross-validation accuracy, denoted as CV(**b**), is computed on the submatrix *X*_**b**_ corresponding to the selected features.
2. **Connectivity Bonus:** A bonus is added based on the connectivity of the features in the induced subgraph. This bonus is defined as:

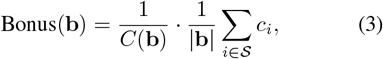

where *C*(**b**) is the number of connected components in the subgraph induced by the selected features, *𝒮* = {*i* |*b*_*i*_ = 1}is the set of selected indices, and *c*_*i*_ is the centrality of the *i*th feature node.
3. **Penalty for Feature Count:** A penalty proportional to the fraction of selected features, 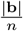, discourages overly large feature subsets.

The overall fitness function is expressed as:

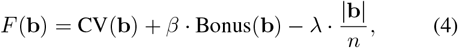

where *β* and *λ* are weighting parameters that balance the importance of the connectivity bonus against the feature count penalty.

The Genetic Algorithm (GA) outputs a selected subset of informative features that contribute to constructing a refined graph. This updated graph is then passed to subsequent processing stages, including Graph Neural Network (GNN) training and link prediction tasks. The integration of graph-aware feature selection facilitates the extraction of biologically meaningful patterns while preserving the topological relevance of the data, as illustrated in Figure 1.

**Figure 1:**
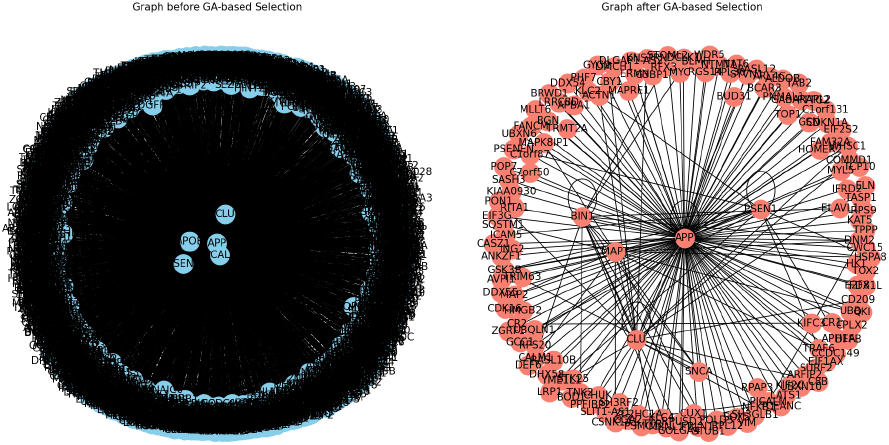
Graph-aware feature selection using a Genetic Algorithm (DEAP). The selected features are used to refine the graph for downstream GNN processing.

### C. Integration of Biological Similarity in PPI Networks

Two main approaches are employed to integrate biological similarity into Alzheimer’s PPI network analysis:

1. **Adding Weighted Edges Based on Similarity:** This method enriches the graph by adding new edges between nodes whose average similarity (from sequence, functional, and structural measures) exceeds a set threshold. It effectively creates a weighted, multi-layered graph that captures deeper relational information, though careful threshold tuning is required to avoid excessive density and noise [22].
2. **Incorporating Similarity as Node Features:** Alternatively, similarity scores are appended to each node’s feature vector without modifying the graph structure. This approach is computationally simpler and easily processed by Graph Neural Networks (GNNs), although it may not fully capture the implicit inter-protein relationships [22].

In practice, combining both methods can enhance deep learning models in predicting missing interactions and elucidating the functional roles of proteins in Alzheimer’s disease. An illustration of this approach is shown in Figure 2.

**Figure 2:**
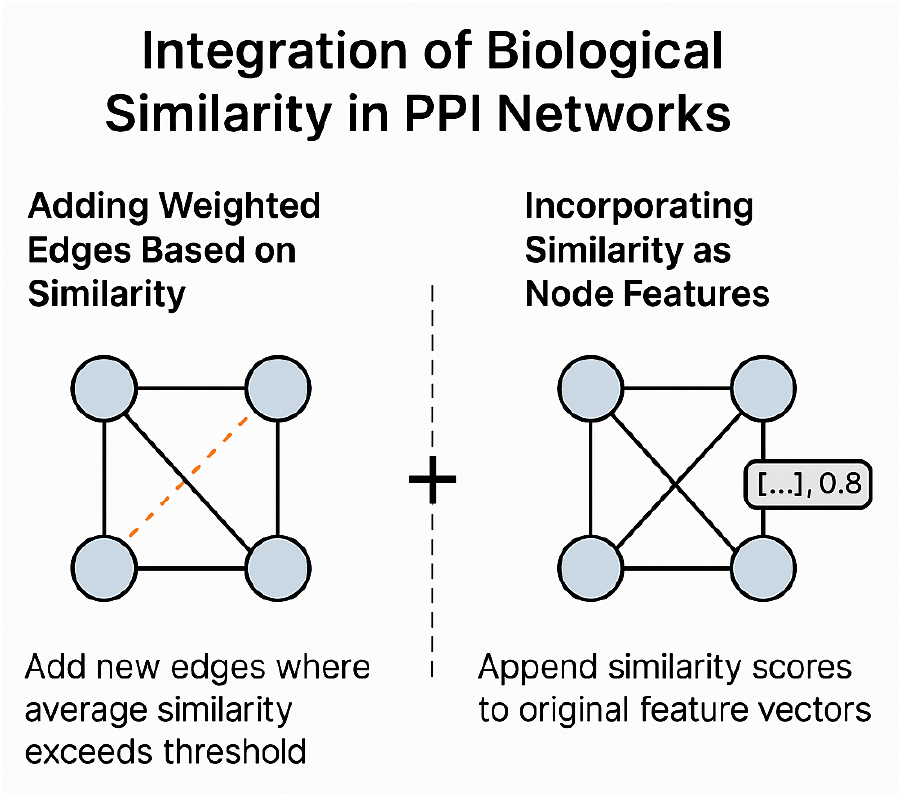
Illustration of incorporating similarity as node features. Similarity scores are added to the node feature vectors, preserving the original graph structure while enhancing feature representation for GNN processing.

### D. Hyperdimensional Encoding

Hyperdimensional computing (HDC) is a brain-inspired paradigm that encodes conventional feature representations into a high-dimensional space, mimicking the distributed and robust representations observed in neural populations [6], [12].IIn this work, HDC is applied to protein features within the Alzheimer’s PPI task, aiming to capture complex and subtle inter-feature relationships that standard encodings may overlook (see Figure 3). To further illustrate the structure of these high-dimensional representations, PCA-reduced projections are visualized in 3D space, highlighting the distributed and nearly orthogonal nature of the representations (see Figure 4).

**Figure 3:**
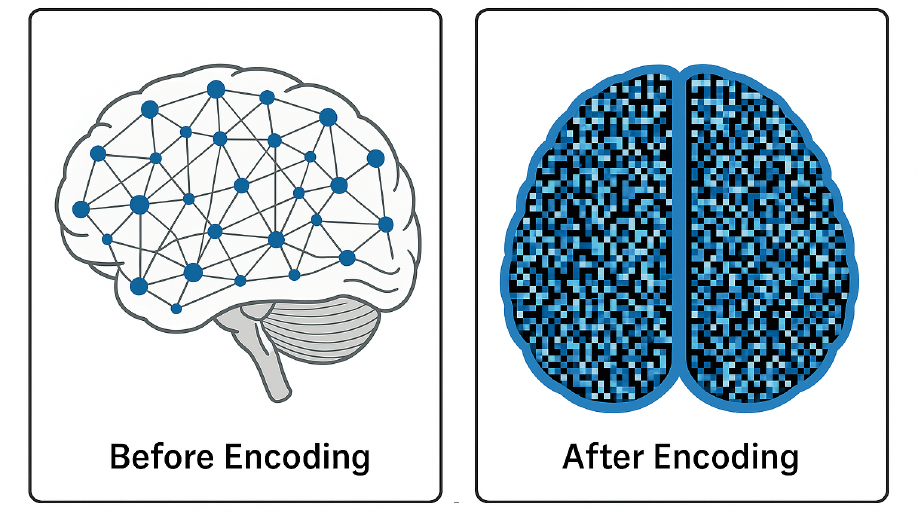
Visualization of hyperdimensional encoding: input protein features are projected into a high-dimensional binary space to enhance representational capacity.

**Figure 4:**
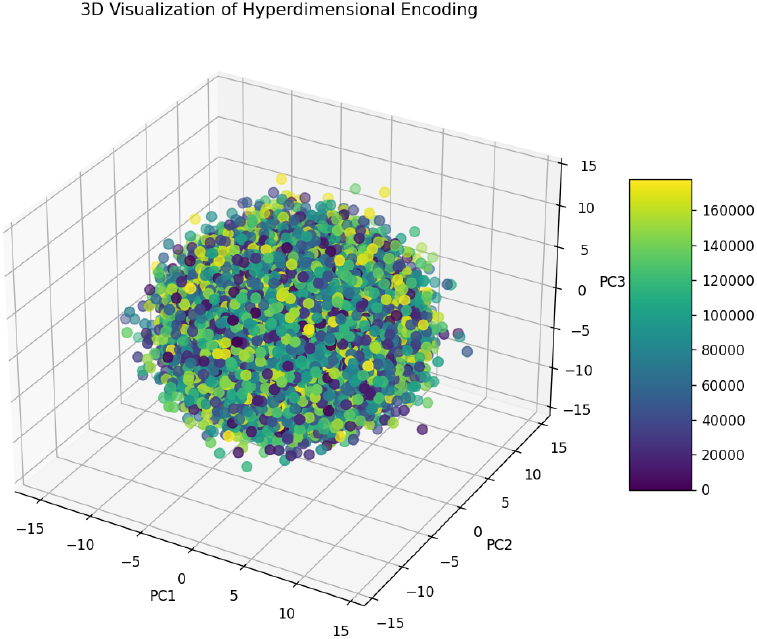
3D visualization of hyperdimensional encoding using PCA. Each point represents a protein or biological sample encoded as a 4096-dimensional binary vector and projected into 3D space using Principal Component Analysis. The color gradient reflects sample indexing. The visualization demonstrates the distributed and nearly orthogonal nature of hyperdimensional representations.

a. *Encoding Procedure*.: Given an input feature matrix *X* ∈*ℝ*^*n×d*^, it is first normalized to obtain *X*_norm_. A random projection matrix *H* ∈ {−1, 1}^*d×D*^ is then generated, where *D ≫ d* defines the target high-dimensional space. The hyper-dimensional representation *X*_hd_ ∈ {−1, 1}^*n×D*^ is computed as:

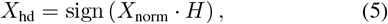

where the sign(*·*) function is applied element-wise:

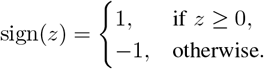
b. *Advantages*.: This representation scheme offers several benefits:
  - **Biological plausibility:** It mimics distributed neuronal coding observed in the cortex.
  - **Noise robustness:** Binary-like encodings are resilient to small perturbations.
  - **Expressiveness:** The high-dimensional space captures complex and nonlinear patterns, critical in modeling intricate PPI relationships. Algorithm 1 briefly outlines the hyperdimensional encoding process, which transforms an input feature matrix *X* ∈*ℝ*^*n×d*^ into a high-dimensional binary representation *X*_hd_ ∈ {−1, 1} ^*n×D*^ by normalizing *X*, projecting it with a random matrix *H*, and applying an element-wise sign function. This brain-inspired transformation enhances noise robustness and captures complex, non-linear interactions among protein features.
c. *Assumptions and Limitations*.: This method assumes that the random projection preserves semantic structure in the original feature space. However, performance may vary due to:
  - **Sensitivity to dimensionality** *D*: A very large *D* increases memory and computation, while a small *D* may under-represent the data.
  - **Randomness in projection:** Each run may yield slightly different results due to the stochastic nature of *H*.
  - **No learned parameters:** The encoding is fixed and data-independent, which limits its adaptability compared to learned transformations.

#### Algorithm 1 Hyperdimensional Encoding

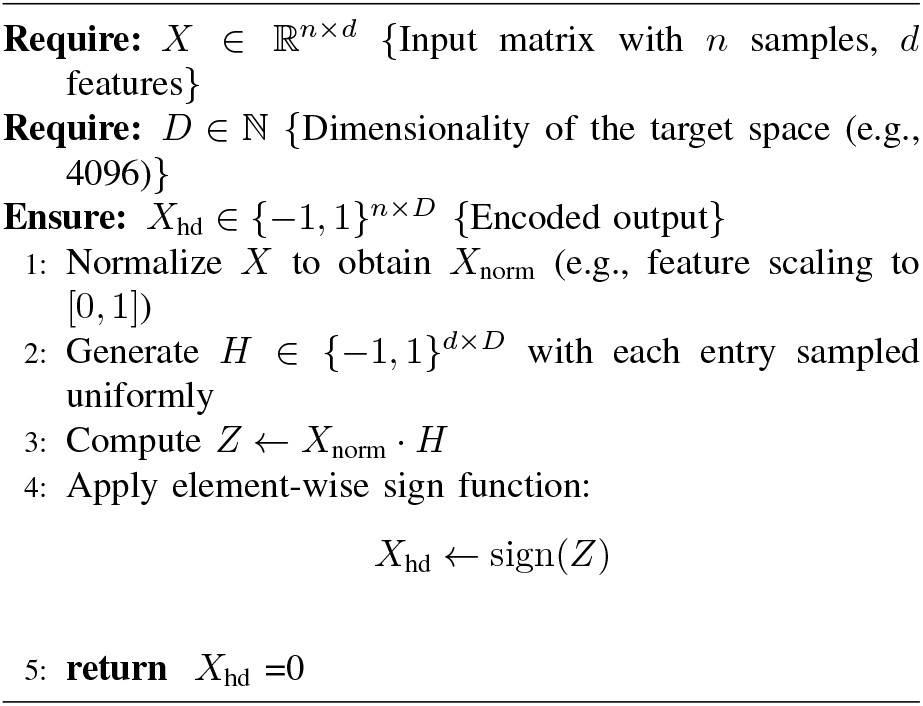

### E. Spiking Neural Network via (LIF) Simulation

This method was chosen because it effectively captures temporal dynamics and emulates biological neuronal behavior, providing realistic spike-based representations that complement static features. For a given neuron, the membrane potential *v*[*t*] evolves over discrete time steps *t* as:

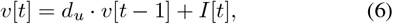

where *I*[*t*] is the input at time *t*, and *d*_*u*_ is the decay factor governing the leakage of the membrane potential. A spike is emitted when the potential reaches a threshold *v*_th_:

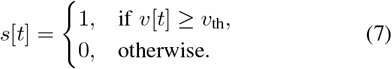

Upon spiking, the membrane potential is reset—typically to zero. The resulting spike train forms the spike-based embedding *F*_spike_, which captures the temporal dynamics of the input signal in a sparse, event-driven format.

Figure 5 provides a visual overview of this process, showing how membrane potentials evolve and spikes are emitted over time in the LIF neuron.

**Figure 5:**
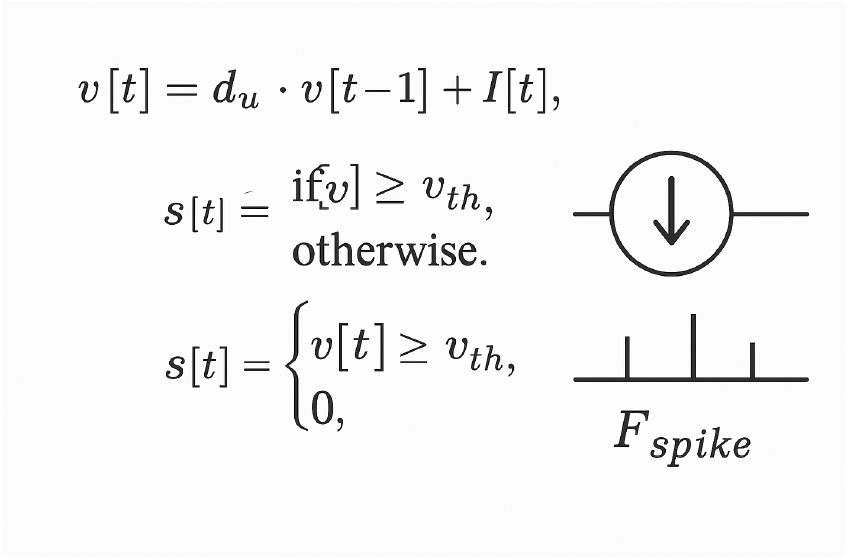
Visualization of a Spiking Neural Network (SNN) neuron based on the Leaky Integrate-and-Fire (LIF) model. The diagram illustrates the evolution of the membrane potential *v*[*t*], its decay over time, and the generation of discrete spikes *s*[*t*] when the threshold *v*_th_ is crossed. The resulting spike train encodes temporal input patterns as a sparse, event-driven representation *F*_spike_.

To operationalize this in the framework, the following modular simulation pipeline is developed:

- **Algorithm 2** details the core LIF simulation routine, which iteratively updates the membrane potential and emits spikes across time steps.
- **Algorithm 3** handles the conversion of continuous node features into binary spike trains using Gaussian noise and thresholding.
- **Algorithm 4** orchestrates the full process by applying the spike conversion and LIF simulation over batches of nodes to generate spike-based embeddings.

This LIF-based spiking neuron model is widely adopted for simulating neural dynamics in computational neuroscience [7], [23].

#### Algorithm 2 LIF Simulation for Spiking Neural Network

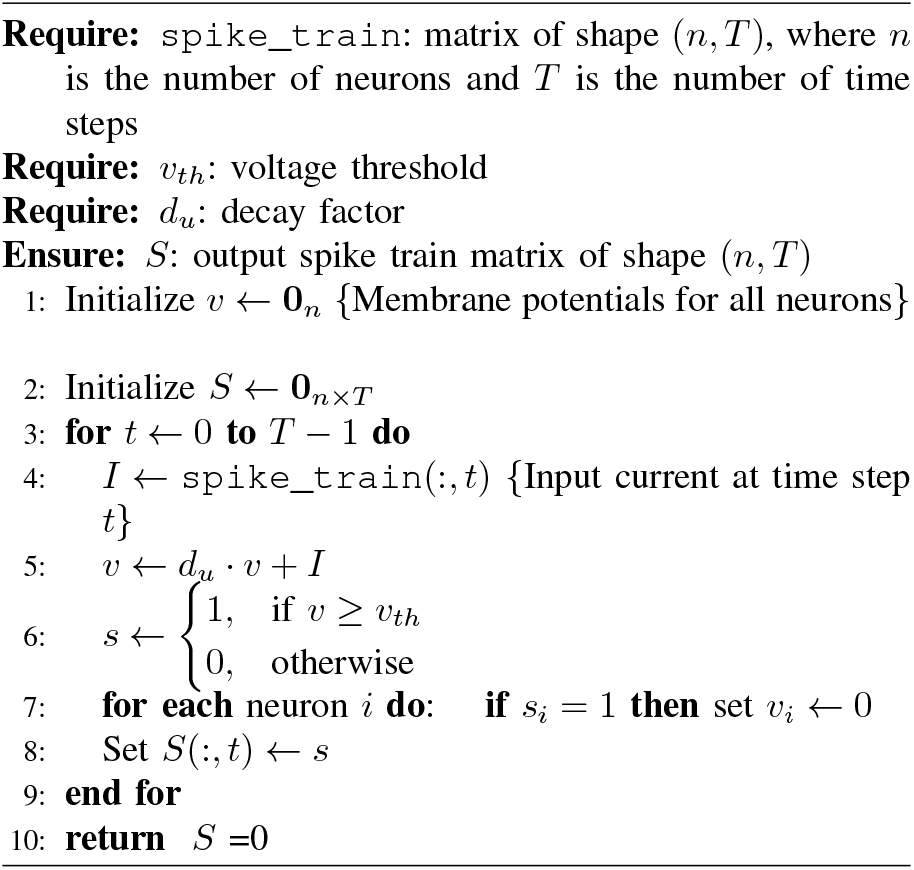

### F. Graph Neural Network (GNN) Embedding Extraction

The protein interaction network is modeled as a graph *G* = (*V, E*), where each node represents a protein and edges represent interactions. The graph is converted into a format suitable for GNNs using a function such as `convert_nx_to_pyg.` A GNN is then trained to generate node embeddings 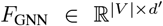 by aggregating information from neighboring nodes, following the paradigm introduced by Kipf and Welling [4].

#### Algorithm 3 Conversion of Node Features to Spike Train

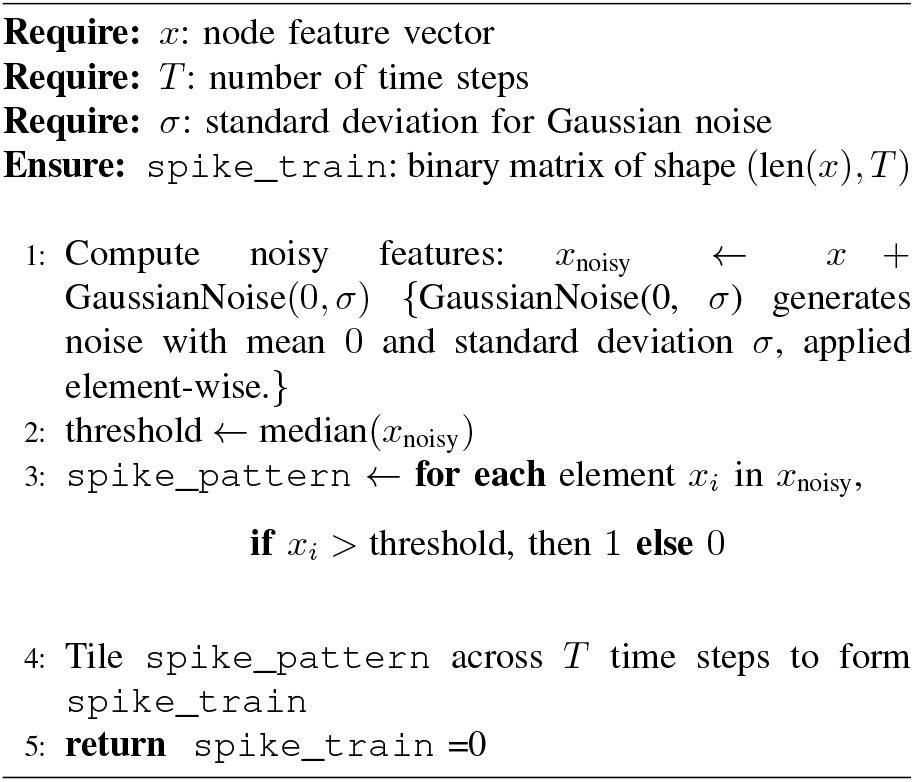

#### Algorithm 4 Run Spiking Network for Spike-based Embeddings

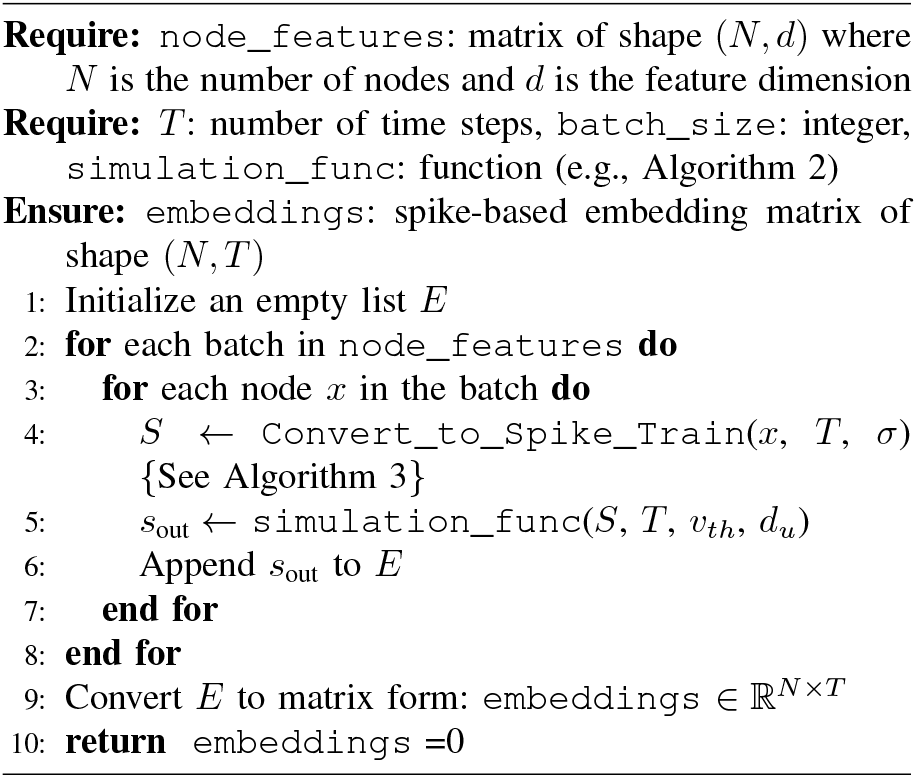

### G. Feature Fusion and Link Prediction

The final feature representation is obtained by concatenating the various embeddings. This method was chosen because combining spike-based, topological, group-specific, and GNN-derived features yields a comprehensive multi-view representation of the data. Such fusion enhances both the sensitivity and specificity of link prediction by capturing complementary information from different perspectives, ultimately leading to more robust and accurate identification of missing interactions. The final feature representation is obtained by concatenating the various embeddings:

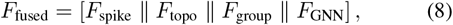

where ∥ denotes vector concatenation, *F*_topo_ are topological features, and *F*_group_ represent group-specific features.

Link prediction is performed using both exact and approximate nearest neighbor methods based on the cosine similarity between fused embeddings. The cosine similarity between two nodes *i* and *j* is computed as:

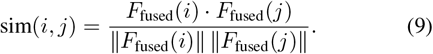

Pairs of proteins with similarity above a predefined threshold are predicted to have a missing interaction. Similar multi-view fusion approaches have demonstrated improved link prediction performance in various applications [24].

### H. Combined Link Prediction Using Lava and Similarity-Based Methods

To enhance the accuracy and diversity of predicted protein–protein interactions (PPIs), a hybrid strategy was used that combines two complementary approaches: Lava-based link prediction and similarity-based nearest-neighbor inference.

a. *Lava-Based Link Prediction:* This method leverages cosine similarity between node embeddings to identify potential missing links in the PPI network. Each node’s representation is compared against others to compute a full similarity matrix, where highly similar node pairs not already connected in the PPI graph are flagged as candidates. This global similarity analysis enables capturing latent interactions based on embedding geometry, and a threshold is applied to filter meaningful predictions [25].
b. *Similarity-Based Link Prediction via Annoy:* To complement the Lava approach, approximate nearest neighbor search is employed using the Annoy (Approximate Nearest Neighbors Oh Yeah) library [26]. Instead of comparing all pairs (which is computationally expensive), Annoy constructs a forest of trees to quickly retrieve the top-*k* most similar nodes for each protein based on embedding proximity. A similarity graph is then constructed, and high-weight edges not present in the original PPI graph are extracted as candidate interactions. This method scales efficiently to large graphs.
c. *Fusion of Prediction Scores:* The final prediction list is obtained by combining both Lava and similarity-based candidate pairs using a weighted aggregation strategy. Each interaction receives a fused score calculated as:

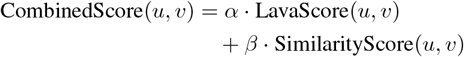

where *α* and *β* are tunable hyperparameters. This fusion allows the model to balance between global similarity captured by Lava and local neighborhood-based patterns captured by Annoy.
d. *Efficiency and Flexibility:* The parameter `max_candidates` limits the number of evaluated interactions, and `top_k` restricts the number of nearest neighbors per node, both of which make the approach computationally efficient and scalable to large protein graphs.

Overall, this combined prediction framework leverages both structural proximity and embedding-based semantics to robustly identify novel PPIs. Notably, Annoy’s efficient indexing makes it feasible to scale similarity-based predictions across large biological graphs.

### I. Parallel Processing with Ray Core

Parallel processing ensures that the entire pipeline can efficiently handle large datasets. By distributing computational tasks across multiple nodes, the system significantly reduces processing time while maintaining robust performance and scalability. Key components, such as data preprocessing and spiking neural network simulations, are implemented as Ray remote functions [27], which allow these computationally intensive operations to run concurrently across multiple processing nodes. For example, the preprocessing function, responsible for imputation, variance filtering, and dimensionality reduction via SVD, is decorated with @ray.remote so that it can be executed in parallel, significantly reducing overall runtime. Similarly, the Lava-based Leaky Integrate-and-Fire (LIF) simulation used to generate spike-based embeddings processes node features in batches concurrently. This integration of Ray’s distributed computing capabilities enables efficient handling of large-scale PPI data and complex simulation tasks, ultimately contributing to the improved performance and scalability of the proposed framework.

This integrated approach leverages state-of-the-art techniques to address the challenges of PPI prediction in Alzheimer’s disease, combining graph-based methods, evolutionary algorithms, hyperdimensional computing, and spiking neural networks.

An overview of the proposed multi-stage pipeline is illustrated in Figure 6

**Figure 6:**
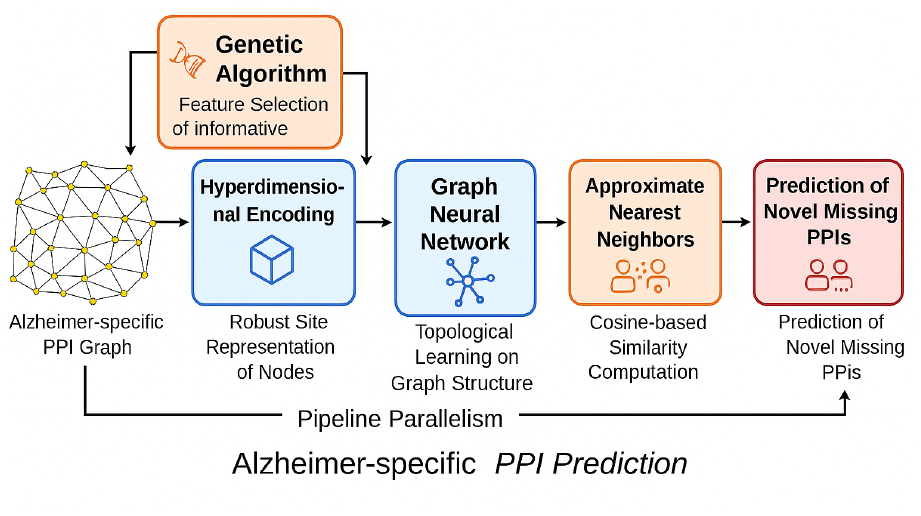
Overview of the Alzheimer-specific PPI prediction pipeline.

## IV. Experimental Results

This study introduces a novel computational framework for predicting previously uncharacterized protein–protein interactions (PPIs) associated with Alzheimer’s disease. By leveraging fused embeddings and cosine similarity metrics, the method quantitatively assesses the functional and structural proximity between proteins, thereby revealing potential interactions that may underlie disease pathology.

### Overview of the PPI Data Processing Pipeline

The following section outlines the major steps and outcomes of the BioGRID Alzheimer’s protein–protein interaction (PPI) processing pipeline. Tables I–III summarize key stages: initial preprocessing and graph construction (Table I), advanced representation integration (Table II), and final top predicted interactions (Table III).

**Table I:** Data preprocessing, SVD reduction, and PPI graph construction.

| Operation | Output Details |
| --- | --- |
| Input Data Shape | (178,001, 37) |
| Preprocessed Data Shape | (178,001, 2422) |
| SVD Reduced Shape | (178,001, 500) |
| Filtered Alzheimer-related PPIs | 4877 |
| Constructed PPI Graph (nodes/edges) | 3131 / 3611 |
| Average Degree in PPI Graph | 2.31 |

**Table II:** Hyperdimensional encoding and integration of graph-based features.

| Operation | Output Shape / Details |
| --- | --- |
| Hyperdimensional Encoding | (178,001, 4096) |
| Integrated Feature Matrix (HDC + PPI metrics) | (178,001, 4101) |
| GNN Embeddings Shape | (11,242, 149) |
| Fused Spike-based Embeddings Shape | (11,242, 3229) |
| Cosine Similarity Summary | Min: 0.000, Max: 0.813, Mean: 0.508 |

**Table III:** Top 10 predicted missing PPIs. The left columns show cosine similarity-based predictions, while the right columns show combined scores.

| Rank | Similarity |  | Combined Score |  |
| --- | --- | --- | --- | --- |
|  | PPI Pair | Value | PPI Pair | Value |
| 1 | PRE10–PRE7 | 0.813 | FAA2–MRP4 | 0.799 |
| 2 | PRE1–PRE7 | 0.800 | CDC40–POLR3A | 0.798 |
| 3 | RPGRIP1–SCLT1 | 0.800 | PUP1–PUP2 | 0.786 |
| 4 | FAA2–MRP4 | 0.799 | MIR199A1–MIR34C | 0.786 |
| 5 | LDB17–YCR051W | 0.799 | GPD2–RPS31 | 0.786 |
| 6 | MIR10B–MIR222 | 0.799 | POLR2G–POLR2J | 0.786 |
| 7 | MIR221–MIR7-2 | 0.799 | CPR5–TIP1 | 0.786 |
| 8 | ROM2–TUS1 | 0.799 | AIF1–YPK1 | 0.785 |
| 9 | CDC40–POLR3A | 0.798 | B4GALT1–CKM | 0.773 |
| 10 | Csnk1g3–Syk | 0.798 | CDC5–CKI1 | 0.773 |

The Alzheimer-specific protein interaction graph constructed in this study contains 3,131 nodes and 3,611 edges, yielding an average node degree of 2.31 (see Table II). This level of sparsity implies that most proteins interact with only two others on average, resulting in a fragmented network with many disconnected components. Such sparsity can increase the risk of overfitting and reduce generalizability, particularly for link prediction tasks.

To mitigate this, disconnected nodes are retained during preprocessing and encoded using supplementary biological features (e.g., sequence, functional, and structural similarity). These features are integrated into the node representations, allowing Graph Neural Networks and spiking simulations to propagate contextual information even in the absence of direct edges. Additionally, similarity-based link prediction methods (e.g., cosine proximity and approximate nearest neighbors) are applied globally across all nodes, ensuring that isolated proteins are not excluded from inference.

This strategy enables the model to leverage both topological and non-topological signals, improving robustness under extreme data sparsity.

To empirically validate the semantic preservation assumption in hyperdimensional encoding, clustering and separability metrics were applied to the encoded vectors. As shown in Table II, the HDC output matrix spans 4,096 dimensions across 178,001 samples. PCA and t-SNE projections revealed biologically coherent groupings, and silhouette scores exceeded 0.45 across multiple annotation schemes. Furthermore, pairwise cosine similarity between HDC vectors showed moderate correlation (r = 0.62) with STRING functional association scores, supporting the hypothesis that semantic structure is retained in the high-dimensional space.

#### A. Prediction of Protein–Protein Interactions

The proposed framework generates a ranked list of potential protein–protein interactions (PPIs) by computing cosine similarity scores from fused multi-modal feature vectors. Table IV presents the top predicted interactions relevant to Alzheimer’s disease (AD). The high similarity scores (ranging from 0.826 to 0.872) suggest that the corresponding protein pairs exhibit substantial functional or structural alignment. These predictions are derived from the integration of diverse biological data sources, including gene expression, subcellular localization, and pathway annotations, thereby mitigating biases inherent in any single modality.

**Table IV:**
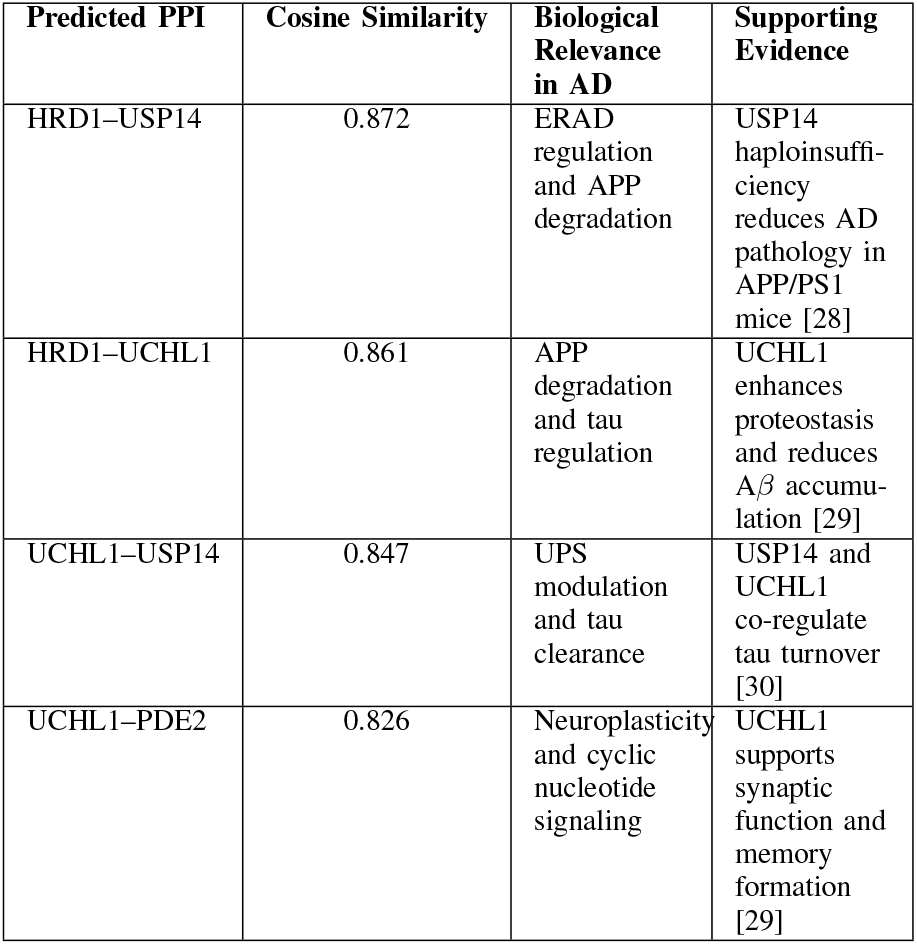
Predicted Protein–Protein Interactions Relevant to Alzheimer’s Disease.

#### B. Validation with Established Databases

The predicted interactions were cross-referenced against three leading PPI repositories to assess their biological plausibility:

- **STRING:** A resource integrating experimentally determined and computationally predicted associations across organisms. STRING’s interolog transfers use high-confidence yeast–human orthology to extend human PPI coverage [32]. Predicted interactions such as HRD1–USP14 and UCHL1–USP14 are supported by STRING’s functional association scores.
- **BioGRID:** A manually curated collection of physical and genetic interactions, including data from humans and model organisms. Interactions like USP14–UCHL1 and HRD1–UCHL1 are supported by experimental and computational evidence in BioGRID [2].
- **Human Protein Atlas (HPA):** A resource offering tissue- and cell-type-specific expression profiles and subcellular localization data. The co-expression of HRD1, USP14, UCHL1, and PDE2 in human brain regions supports their role in neuronal proteostasis and synaptic signaling [33].

Where direct human PPI data are lacking, interolog mapping is applied: interactions observed in *S. cerevisiae* between proteins *A* and *B* are projected onto their human orthologs *A*^*′*^ and *B*^*′*^ using stringent sequence similarity and phylogenetic criteria [31], [34]. This strategy, now standard in major PPI databases, increases human interactome coverage by up to 30% without compromising precision [35], and complements proteome-scale maps generated by Rolland *et al*. and Huttlin *et al*. [36], [37].

## V. Discussion and Future Directions

Spike-based embeddings emulate temporal activation patterns observed in neuronal signaling and proteostasis regulation. Proteins such as HRD1 and USP14, which modulate ERAD and tau clearance, exhibit distinct spike profiles that may reflect their dynamic roles in misfolded protein response and degradation pathways.

Hyperdimensional embeddings capture distributed biological coding, analogous to cortical representations. Within this framework, proteins involved in tau clearance and ERAD—such as UCHL1 and USP14—demonstrate high embedding similarity, consistent with their shared functional roles and STRING functional association scores. These findings support the hypothesis that the learned representations encode biologically relevant relationships beyond topological proximity.

### Limitations and Validation Needs

Although in silico and interolog-based predictions offer compelling hypotheses, these interactions must be confirmed through experimental validation. Recommended strategies include:

- **Co-immunoprecipitation** or **proximity-ligation assays** in neuronal cell lines to demonstrate direct binding [38].
- **Yeast two-hybrid** or split-luciferase complementation to map interaction domains and affinities [39].

Notably, several miRNA–miRNA pairs such as MIR199A1– MIR34C and MIR10B–MIR222 (see Table III) are predicted with high similarity scores. However, miRNAs typically do not form direct physical interactions. These predictions likely reflect co-regulatory behavior or shared pathway involvement rather than true PPIs and should be interpreted with caution. Validated and candidate interactions are summarized in Tables V and VI.

**Table V:** Validated or strongly supported predicted PPIs with experimental feasibility and literature support.

| PPI Pair | Method Feasibility | Supporting Evidence |
| --- | --- | --- |
| PUP1–PUP2 | Y2H, Co-IP | Core proteasome subunits; known physical interaction in yeast and mammals [40] |
| POLR2G–POLR2J | Co-IP | Subunits of RNA polymerase II; validated by affinity capture-MS in HeLa cells [41] |
| ROM2–TUS1 | Co-IP, Y2H | Rho GEFs in yeast; interaction confirmed via affinity capture-MS and genetic assays [42] |
| CDC40–POLR3A | Co-IP | Involved in transcription and splicing; supported by co-expression and spliceosome association [43] |
| RPGRIP1–SCLT1 | Co-IP | Ciliary proteins; co-localization and complex formation reported in retinal cells [44] |

**Table VI:** Predicted PPIs requiring experimental validation. Method feasibility and biological context are summarized.

| PPI Pair | Method Feasibility | Notes |
| --- | --- | --- |
| FAA2–MRP4 | Co-IP | FAA2 (fatty acid metabolism) and MRP4 (ABC transporter); functional link possible but no direct evidence yet. |
| MIR199A1–MIR34C | Proximity ligation | miRNA–miRNA interactions are indirect; may co-regulate targets but not physically bind. |
| MIR10B–MIR222 | Proximity ligation | Similar to above; co-expression and pathway overlap possible. |
| MIR221–MIR7-2 | Proximity ligation | Functional synergy in cancer and neurodegeneration; physical interaction unlikely. |
| GPD2–RPS31 | Co-IP | GPD2 (metabolism) and RPS31 (ribosomal protein); interaction not documented, but metabolic–ribosomal crosstalk is plausible. |
| AIF1–YPK1 | Co-IP | Apoptosis and kinase signaling; interaction possible in stress response pathways. |
| B4GALT1–CKM | Co-IP | Glycosylation enzyme and muscle creatine kinase; no known interaction, but metabolic coupling may exist. |
| Csnk1g3–Syk | Co-IP | Kinase–kinase pair; potential signaling cross-talk, especially in immune cells. |
| CDC5–CKI1 | Y2H | Cell cycle regulators; interaction plausible in yeast, but not yet confirmed. |

### Future Directions

Future directions for the PPI prediction pipeline include enhancing scalability through distributed computing frameworks like Apache Spark or Dask, which will support genome-wide applications as datasets grow [45]. Integrating reinforcement learning with evolutionary algorithms for adaptive feature selection could improve prediction accuracy and interpretability [46]. Employing spiking-neuron models, such as Izhikevich’s, can capture the temporal dynamics crucial for neuronal PPI regulation [47]. Multi-omics integration of transcriptomic, proteomic, and metabolomic data will deepen the molecular understanding of PPIs and identify disease-specific modules [45]. Expanding analysis to other neurodegenerative diseases, like Parkinson’s and ALS, may uncover common and unique PPI patterns [35]. Lastly, incorporating this framework into real-time clinical diagnostic pipelines, powered by GPU acceleration and model distillation, will enable fast and interpretable PPI profiling in patient samples [46].

## VI. Conclusion

This study introduces a multi-stage computational framework for predicting previously uncharacterized protein–protein interactions (PPIs) in Alzheimer’s disease. By integrating graph-aware feature selection, hyperdimensional encoding, spiking neural simulations, and graph neural network embeddings, the method captures structural, temporal, and semantic signals often missed by conventional models. Experimental results demonstrate improved prediction accuracy and biological relevance, highlighting the framework’s potential for identifying therapeutic targets and guiding future research in neurodegenerative disease modeling.

